# The temporal paradox of Hebbian learning and homeostatic plasticity

**DOI:** 10.1101/116400

**Authors:** Friedemann Zenke, Wulfram Gerstner, Surya Ganguli

**Affiliations:** Department of Applied Physics, Stanford University, Stanford, CA 94305, USA; Brain Mind Institute, School of Life Sciences and School of Computer and Communication Sciences, Ecole Polytechnique Fédérale de Lausanne, CH-1015 Lausanne EPFL, Switzerland

**Keywords:** synaptic plasticity, Hebbian plasticity, homeostatic plasticity, rapid compensatory processes

## Abstract

Hebbian plasticity, a synaptic mechanism which detects and amplifies co-activity between neurons, is considered a key ingredient underlying learning and memory in the brain. However, Hebbian plasticity alone is unstable, leading to runaway neuronal activity, and therefore requires stabilization by additional compensatory processes. Traditionally, a diversity of homeostatic plasticity phenomena found in neural circuits are thought to play this role. However, recent modelling work suggests that the slow evolution of homeostatic plasticity, as observed in experiments, is insufficient to prevent instabilities originating from Hebbian plasticity. To remedy this situation, we suggest that homeostatic plasticity is complemented by additional rapid compensatory processes, which rapidly stabilize neuronal activity on short timescales.

## Introduction

More than half a century ago, Donald Hebb [1] laid down an enticing framework for the neurobiological basis of learning, which can be succinctly summarized in the well-known mantra, “neurons that fire together wire together” [2]. However, such dynamics suffers from two inherent problems. First, Hebbian learning exhibits a positive feedback instability: those neurons that wire together will fire together more, leading to even stronger connectivity. Second, such dynamics alone would lead to all neurons in a recurrent circuit wiring together, precluding the possibility of rich patterns of variation in synaptic strength that can encode, through learning, the rich structure of experience. Two fundamental ingredients required to solve these problems are stabilization [3], which prevents runaway neural activity, and competition [4–6], in which the strengthening of a synapse may come at the expense of the weakening of others.

In theoretical models, competition and stability are often achieved by augmenting Hebbian plasticity with additional constraints [3, 5, 7]. Such constraints are typically implemented by imposing upper limits on individual synaptic strengths, and by enforcing some constraint on biophysical variables, for example, the total synaptic strength or average neuronal activity [6–12]. In neurobiology, forms of plasticity exist which seemingly enforce such limits or constraints through synaptic scaling in response to firing rate perturbations [13, 14], or through stabilizing adjustments of the properties of plasticity in response to the recent synaptic history, a phenomenon known as homeostatic metaplasticity [6, 11, 15, 16]. Overall, synaptic scaling and metaplasticity, as special cases of homeostatic mechanisms that operate over diverse spatiotemporal scales across neurobiology [17–22], are considered key ingredients that contribute both stability and competition to Hebbian plasticity by directly affecting the fate of synaptic strength.

The defining characteristic of homeostatic plasticity is that it drives synaptic strengths so as to ensure a homeostatic set point [23, 24], such as a specific neuronal firing rate or membrane potential. However, it is important that this constraint is implemented only on average, over long timescales, thereby allowing neuronal activity to fluctuate on shorter timescales, so that these neuronal activity fluctuations, which drive learning through Hebbian plasticity, can indeed reflect the structure of ongoing experience. This requisite separation of timescales is indeed observed experimentally; forms of Hebbian plasticity can be induced on the timescale of seconds to minutes [25–28], whilst most forms of homeostatic synaptic plasticity operate over hours or days [14, 24, 29]. This separation of timescales, however, raises a temporal paradox: homeostatic plasticity then may become too slow to stabilize the fast positive feedback instability of Hebbian learning. Indeed modeling studies that have attempted to use homeostatic plasticity mechanisms to stabilize Hebbian learning [11, 30–34] were typically required to speed up homeostatic plasticity to timescales that are orders of magnitude faster than those observed in experiments (Fig. 1).

This temporal paradox could have two potential resolutions. First, the timescale of Hebbian plasticity, as captured by recent plasticity models fit directly to data from slice experiments [28, 35–38], may overestimate the rate of plasticity that actually occurs *in-vivo*. This overestimate could arise from differences in slice and *in-vivo* preps, or because complex nonlinear synaptic dynamics, both present in biological synapses and useful in learning and memory [39–41], are missing in most, but not all [42–44], data-driven models. While slow plasticity may be a realistic possibility in cortical areas exhibiting plastic changes over days [45, 46], it may not be a realistic resolution in other areas, like the hippocampus, which must rapidly encode new episodic information [47, 48]. The second potential resolution to the paradoxical separation of timescales between Hebbian and homeostatic plasticity may be the existence of as yet unidentified rapid compensatory processes (RCPs) that stabilize Hebbian learning. Below, we explore both the theoretical utility and potential neurobiological instantiations of these putative RCPs.

**Figure 1:**
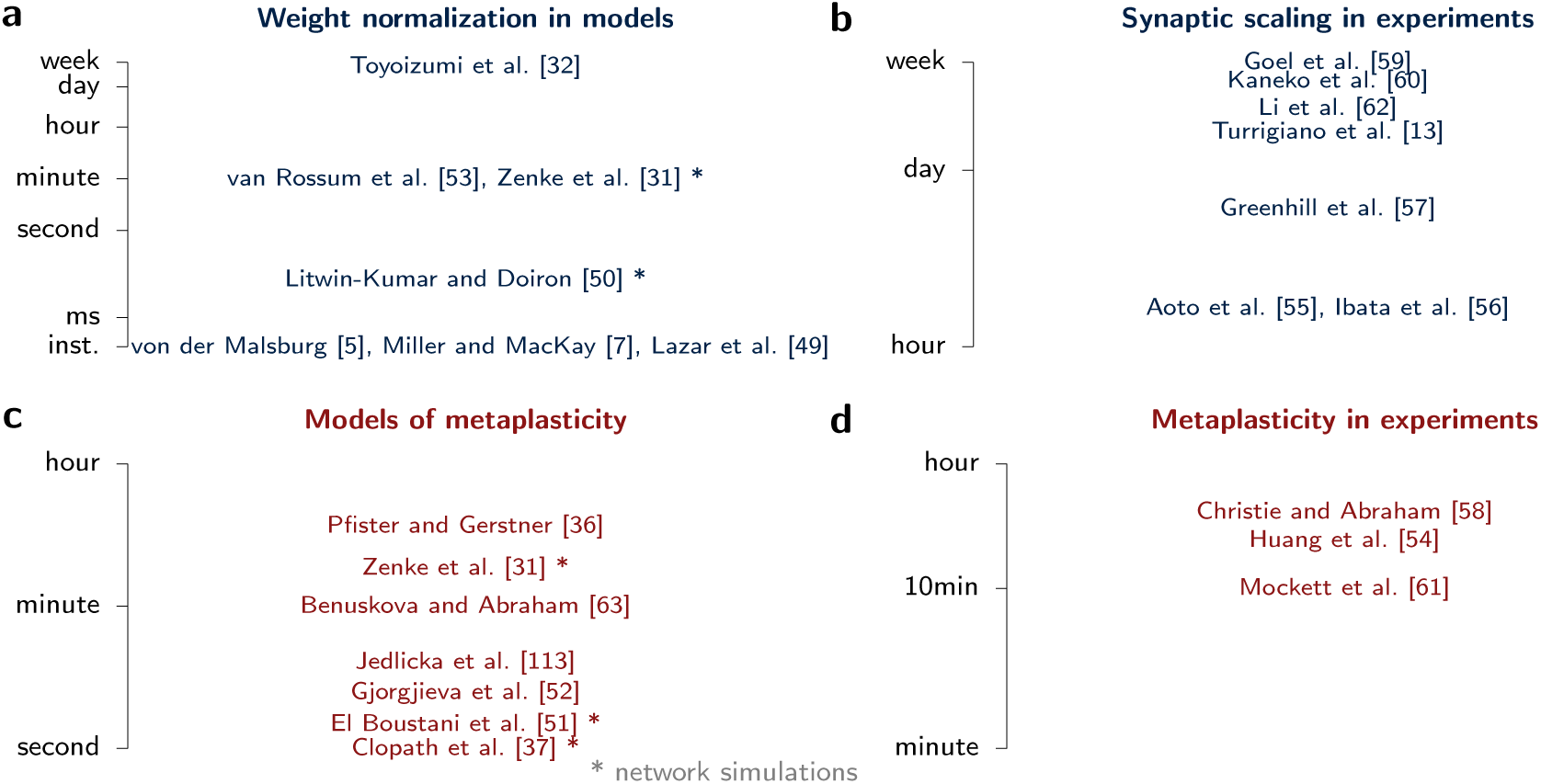
Comparison of the typical timescale of different forms of homeostatic plasticity in models and in experiments. (a) Weight normalization in models. Here we plot the characteristic timescale on which synaptic weights are either normalized or scaled. (b) Synaptic scaling in experiments. Here we plot the typical time at which synaptic scaling is observed. (c) Models of metaplasticity. We plot the characteristic timescale on which the learning rule changes. (d) Metaplasticity in experiments. Here we show the typical timescale at which metaplasticity is observed.

### The temporal paradox of Hebbian and homeostatic plasticity

To understand the theoretical necessity for RCPs to stabilize Hebbian plasticity, it is useful to view a diversity of synaptic learning models through the unifying lens of control theory (Fig. 2a). Here we can view the “fire together, wire together” interplay of neuronal activity and Hebbian synaptic plasticity as an unstable dynamical system. Also, we can view a compensatory process as a feedback controller that observes some aspect of either neuronal activity or synaptic strength, and uses this observation to compute a feedback control signal which then directly affects synaptic strength so as to stabilize global circuit activity. Indeed homeostatic plasticity is often thought of as a negative feedback control process [23, 24, 64, 65]. In general, the delay in any feedback control loop must be fast relative to the time-scale over which the unstable system exhibits run-away activity [66]. If the loop is slightly slow, the run-away will start before the stabilizing feedback arrives, generating oscillations within the system. In some cases, if the loop is even slower, the unstable runaway process might escape before stabilization is even possible (Fig. 2b).

Such oscillations and run-away are demonstrated in Fig. 3 for several compensatory mechanisms with feedback timescales that are chosen to be too slow, including the Bienenstock-Cooper-Munro (BCM) rule [6, 67], and triplet spike-timing-dependent plasticity (STDP) [36] with either synaptic scaling [13, 53] or a metaplastic sliding threshold, as stabilizing controllers. For example, the BCM rule (Fig. 3a) can be thought of as a feedback control system where the controller observes a recent average of the postsynaptic output firing rate of a neuron, and uses this information to control both the sign and amplitude of associative plasticity; if the recent average is high (low), plasticity is modulated to be anti-Hebbian (Hebbian). However, to achieve stability, the BCM controller must average recent output-activity over a short enough time-scale to modulate plasticity before this activity itself runs away (Fig. 3a; [11, 31, 32]). This result is not limited to BCM like rate models, but applies equally to STDP models which rely on similar metaplasticity processes to ensure stability (Fig. 3bc; [31, 36, 37, 52]). These modified STDP rules can be thought of as employing feedback controllers which also observe a recent average of output firing rate, but use this information to modulate the STDP window, thereby stabilizing the system by changing relative rates of long-term potentiation (LTP) and long-term depression (LTD).

**Figure 2:**
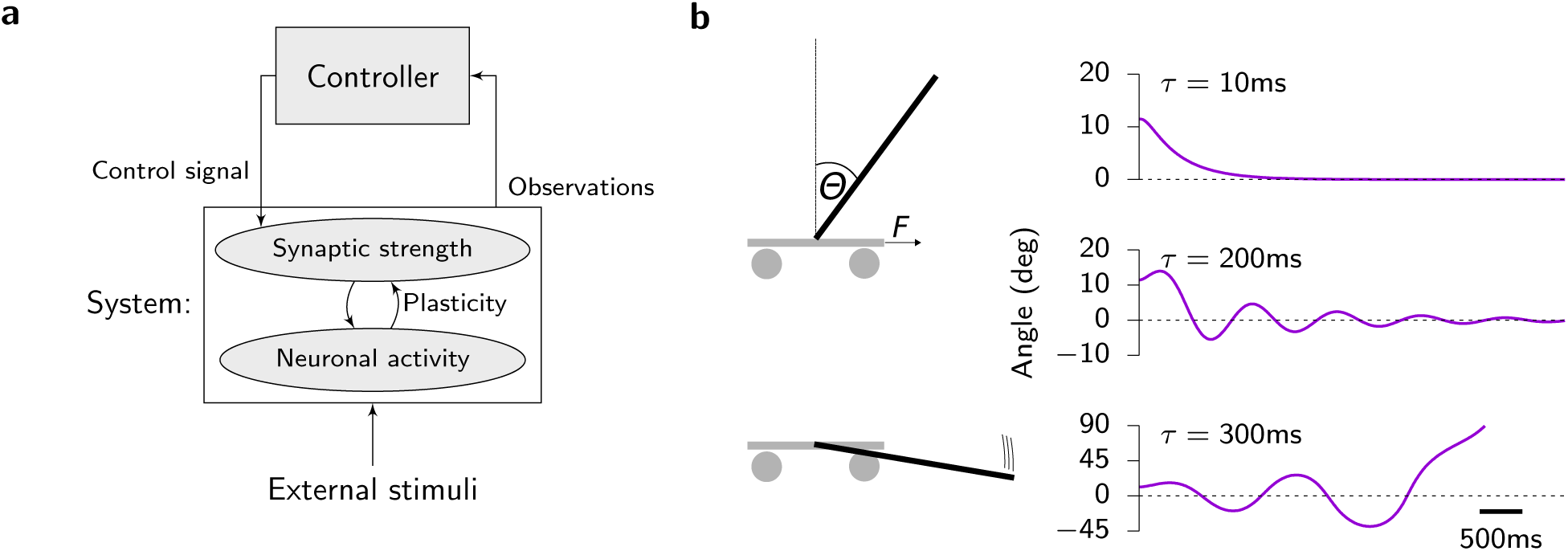
A control theoretic view of homeostatic plasticity and Hebbian learning. (a) The coupled dynamics of Hebbian plasticity and neural activity constitutes an unstable dynamical system. Homeostatic plasticity, and more generally any compensatory mechanism, can be viewed as a controller that observes aspects of neuronal activity and synaptic strengths, and uses these observations to compute a feedback control signal that acts on synaptic dynamics so as to stabilize circuit properties. (b) The cart pole problem is a simple example of stabilizing a non-linear dynamical system with feedback. The task of the controller is to exert horizontal forces on the cart to maintain the rod (*m* = 1kg) in an upright position. For simplicity we assume a weightless cart with no spatial constraints on the length of the track. The controller has access to an exponential average (time constant *τ*) over recent observations of both the angle *θ* and angular velocity *θ* of the pole. For small values of *τ* the controller can successfully maintain the rod upright (*θ* ≈ 0; top panel). However, as *τ* and the associated time lag in the observed quantities gets larger, oscillations arise (middle panel) and eventually the system becomes unstable (bottom panel).

Another class of models relies on re-normalization of afferent synaptic weights to stabilize Hebbian plasticity. We distinguish between models with instantaneous algorithmic re-normalization of the weights [4, 5, 7, 50] and models which proportionally scale synaptic weights in an activity dependent way [8, 31, 68]. In the former case the controller observes total synaptic strength and uses this information to adjust synapses to keep this total strength constant. In the latter case, the controller observes neuronal firing rate, or recent average thereof, and uses this information to proportionally adjust afferent synaptic weights to enforce either a specified target output, or a total synaptic strength. Just as in the case of metaplasticity discussed above, the temporal average of the activity sensor has to be computed over a short timescale, related to the timescale over synaptic strengths and neuronal activity change, to ensure stability (Fig. 3d [31–33]). Moreover, the rate at which the synaptic scaling process itself causes synaptic strengths to scale must be finely tuned to a narrow parameter regime that is neither too fast nor too slow [31–33, 53]; if too slow, then stabilization is not possible, while if too fast, the stabilization process overshoots, causing oscillations.

Finally, some STDP models can be intrinsically stable, especially in feedforward circuits. One example are pair-based STDP models in which the integral of the STDP window is slightly biased toward depression. When weights are additionally limited by hard bounds, this can lead to bimodal weight distributions and firing rate stabilization [9, 69]. However, these learning rules typically require fine-tuning and become unstable when input correlations are non-negligible. Other stable models arise from a weight dependence in the learning rule such that high (low) synaptic strength makes LTP (LTD) weaker [53, 70–74]. Such a stabilizing weight dependence is advantageous because it can lead to more plausible unimodal weight distributions as observed empirically [72, 74]. However, the unimodal weight distribution, by precluding multi-stability in the configurations of synaptic strengths, typically leads to the rapid erasure of synaptic memory traces in the presence of background activity driving plasticity [73, 75]. Moreover, when such weight dependent synaptic learning rules are embedded in a recurrent neuronal circuit without any additional control mechanisms, they can succumb to runaway neural activity as experience dependent neural correlations emerge in the recurrent circuit [73, 76, 77].

**Figure 3:**
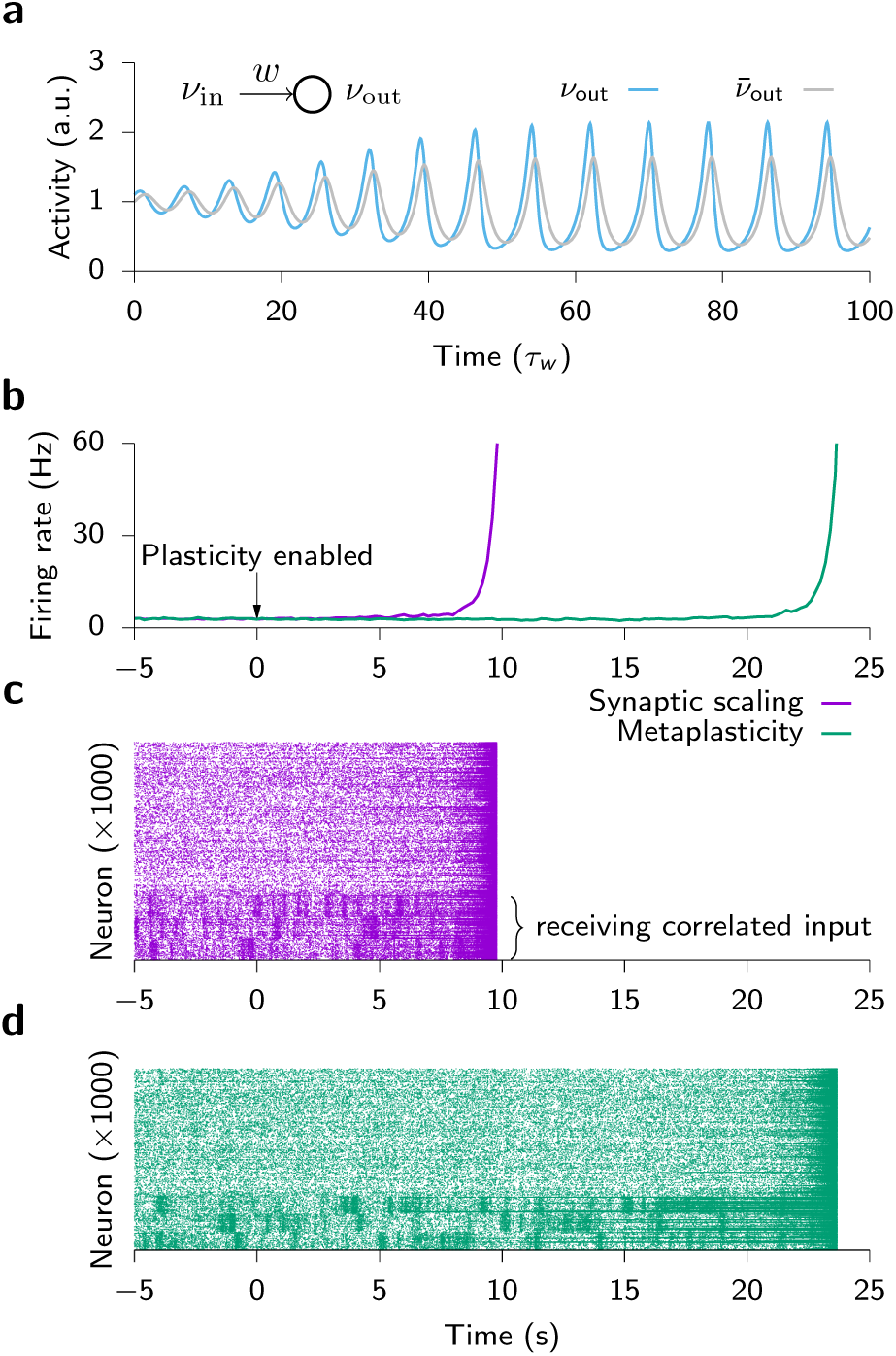
Instability in different plasticity models. (a) Unstable oscillations in the BCM model for a simple feed-forward circuit [6, 11]. Model: 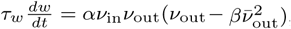, 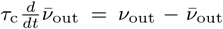 with 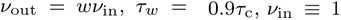 in arbitrary units (a.u.) and α and β are dimensionful scalar constants that ensure correct units; we take simply α = β = 1. Here 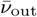 can be thought of as the observation of a controller, corresponding to an average of output neuronal activity *ν*_out_ over time-scale *τ*_c_. Moreover, the multiplicative term in parenthesis in the weight dynamics can be interpreted as a control signal that modulates both the sign and amplitude of associative plasticity, dictating a stabilizing anti-Hebbian rule if the recent average 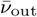 is too large. If the control dynamics T_c_ is too slow relative to the synaptic plasticity dynamics *τ*_*w*_, unstable oscillations arise. (b–d) Runaway activity in a recurrent neural network simulation consisting of 25000 excitatory and inhibitory integrate-and-fire neurons and plastic excitatory synapses using a minimal triplet STDP model [36] with different homeostatic mechanisms such as sliding threshold metaplasticity (violet) and synaptic scaling (green), both described in detail in [31]. (b) Population firing rate as a function of time. (c,d) Raster plot of spiking activity. The bottom 3 groups of 100 neurons received rate modulated spiking input with 100ms correlation time constant to emulate sensory input to a small set of neurons. The timescale of sliding threshold metaplasticity was *τ*_c_ = 3mins. The timescale for the rate detector and the scaling dynamics for synaptic scaling were *τ*_c_ = 10s and *τ*_scl_ = 1h respectively. Because of these slow stabilization dynamics, the fast interplay between Hebbian plasticity and recurrent network dynamics leads to rapid population firing rate destabilization within 10 to 20 seconds for both learning rules.

In summary, empirical findings and control-theoretic considerations suggest that compensatory mechanisms capable of stabilizing Hebbian plasticity must operate on tightly constrained timescales. In practice, such compensatory mechanisms must act on similar or even faster timescales than Hebbian plasticity itself [31, 32, 36, 37, 50, 52, 68]. STDP, as one of the most common manifestations of Hebbian plasticity in the brain, can be induced in a matter of seconds to minutes [25–27, 78, 79]. Homeostatic plasticity, on the other hand, acts on much longer timescales of hours to days [14, 16, 19, 29, 55, 56, 80]. This separation of timescales poses a temporal paradox as it renders most data-driven STDP models unstable [3, 81]. In spiking network models, this instability has severe consequences (Fig. 3b–d); it precludes the emergence of stable synaptic structures or memory engrams [31, 73, 76], unless the underlying plasticity models are augmented by RCPs [31, 50, 51, 68, 82]. Overall, these considerations suggest that one or more RCPs exist in neurobiological systems, which are missing in current plasticity models.

### Putative rapid compensatory processes

What putative RCPs could augment Hebbian plasticity with the requisite stability and competition? Here we focus on several possibilities, operating at either the network, the neuronal or the dendritic level (Fig. 4a). However, we note that these possibilities are by no means exhaustive.

At the network level, recurrent or feedforward synaptic inhibition could influence and potentially stabilize plasticity at excitatory synapses directly. For instance, [83] demonstrated via dynamic-clamp that a simulated increase in total excitatory and inhibitory background conductance, which could originate from elevated levels of network activity, rapidly reduces the amplitude of LTP, but not LTD, of the STDP window. This rapid effect may be mediated through changes in calcium dynamics in dendritic spines and could constitute an RCP. Another study [84] showed that global inhibition in a rate-based network model is sufficient to stabilize plasticity at excitatory synapses with a sliding presynaptic and fixed postsynaptic plasticity threshold. Finally, using a model-based approach, Wilmes et al. [85] have proposed that dendritic inhibition could exert binary switch-like control over plasticity by gating back-propagating action potentials.

Other modeling studies have suggested a role for inhibitory synaptic plasticity (ISP) [86, 87], instead of non-plastic inhibition, in stabilizing Hebbian plasticity. It has been suggested, for instance, that ISP in conjunction with a fixed plasticity threshold at the excitatory synapse could have a similar effect as the sliding threshold in the BCM model [88]. Finally, in some experiments excitatory and inhibitory plasticity are not integrated as independent events, but can influence each other. For instance, there are cases in which induction of ISP alone can flip the sign of subsequent plasticity at excitatory synapses [89]. While inhibition and ISP may act as RCPs, a clear picture of how these elements tie together has not yet emerged. Further experimental and theoretical work is required to understand their potential for acting as RCPs.

**Figure 4:**
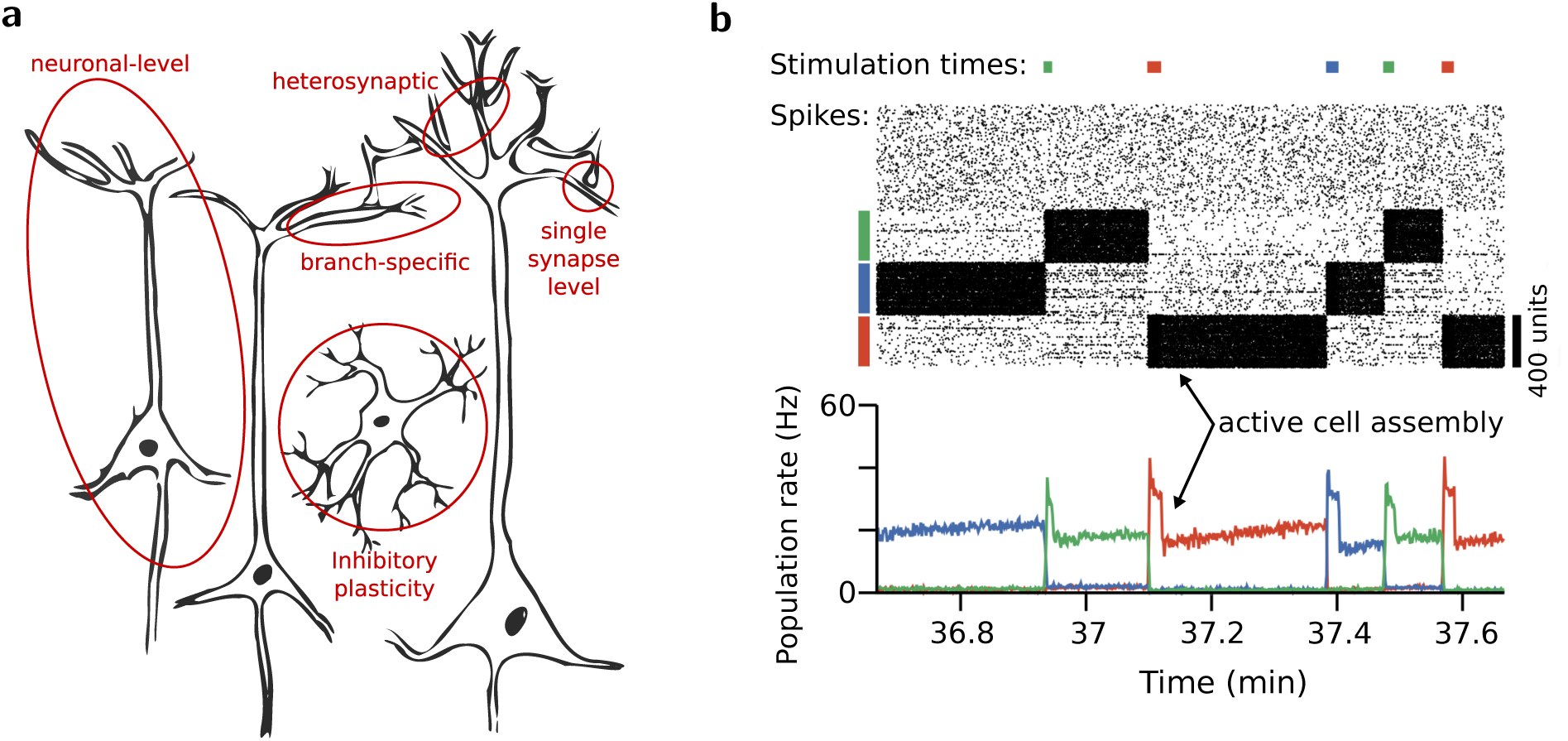
(a) Schematic representation of the many potential loci of rapid compensatory processes. (b) Example of a plastic network model displaying attractor dynamics in three distinct cell assemblies. Because these dynamics outlast the stimulus that triggers them they have been considered as neural substrate for working memory. The top panel shows a spike raster, whereas the bottom panel shows the population firing rates of the three cell assemblies. This particular model uses an augmented triplet STDP learning rule (cf. Fig. 3) in which heterosynaptic plasticity and single-synapse-level plasticity have been added as stabilizing RCPs. Moreover, this model relies on a slow form of inhibitory plasticity which normalizes the overall network activity and ensures that cell assemblies do not grow indefinitely over time by recruiting additional neurons. This provides a proof of principle that a biologically inspired learning rule can indeed be stabilized by a sensible combination of RCPs, whereas the same learning rule endowed with slow compensatory mechanisms leads to run-away dynamics (cf. Fig. 3b–d). Figure adapted from Zenke et al. [82].

Neuromodulation may also play an important role in stabilizing plasticity. Neuromodulators have been implicated in both homeostatic signalling [21] and in gating the expression of synaptic plasticity [90-93]. However, to successfully serve as RCP, neuromodulatory mechanisms have to either drive rapid compensatory changes directly, or substantially reduce the average rate of Hebbian plasticity *in-vivo* to enable slower forms of homeostatic plasticity to preserve stability. While a detailed account of the role of neuromodulators goes beyond the scope of this article (but see [93, 94] for reviews), here we note that several neuromodulators at least partially meet one of these core requirements for RCPs. For example, nitric oxide (NO), which is involved in synaptic homeostasis and plasticity [95], is released in response to increased NMDA activity and decays within some tens of seconds [96]. Because cell membranes are permeable to NO, the molecule diffuses rapidly and thus could potentially act as a fast proxy of bulk neuronal activity that can be read out locally [97]. Similarly, dopaminergic transmission can be fast [98] and is known to affect induction, consolidation and possibly maintenance of synaptic long-term plasticity [91, 93, 99–101]. A sensible gating strategy implemented by dopamine or other neuromodulators could result in a drastically reduced and more manageable average rate of plasticity *in vivo*. In contrast, *in vitro* such neuromodulatory mechanisms might be disengaged, thus potentially creating less natural and much faster plasticity rates compared to *in vivo*. Apart from the neuromodulatory system, recent work has highlighted the complexity of the local interactions between astrocytes and synaptic plasticity which may also act as RCPs [102–104]. However, to unequivocally answer which aspects of neuromodulation and glial interactions constitute suitable RCPs will require further experimental and theoretical work.

A more well studied possibility for an RCP is heterosynaptic plasticity, which operates at the level of individual neurons or potentially dendritic branches. Heterosynaptic plasticity refers to a non input-specific change at *other* synapses onto a neuron that are not directly activated ([30, 105–108]). Its viability as putative RCP arises from the fact that some forms of heterosynaptic plasticity can be induced rapidly, and moreover, similar to synaptic scaling, can show aspects of weight normalization [106]. For instance, a rapid form of heterosynaptic plasticity, in which neuronal bursting causes bi-directional weight-dependent changes in afferent synapses, has been observed recently [107, 109]. While the observed weight-dependence is reminiscent of Oja’s rule [8], as strong synapses weaken, it is not identical because weak synapses can also strengthen. Nevertheless, this form of heterosynaptic plasticity has been shown to prevent runaway of LTP in models of feedforward circuits [30] and has been demonstrated to co-occur with Hebbian plasticity in experiments [109].

Recently, the utility in stabilizing runaway LTP has also been demonstrated in a recurrent network model of spiking neurons [82], in which a similarly burst dependent form of heterosynaptic plasticity is crucial to ensure stable formation and recall of Hebbian cell assemblies (Fig. 4b). Moreover, the work suggests a potential role of heterosynaptic plasticity in triggering the reversal of LTP and LTD [110].

At the level of dendritic branches, a recently described form of structural heterosynaptic plasticity [108], involves local dendritic competition between synapses. Specifically, glutamate induced structural synaptic potentiation of a set of clustered dendritic spines causes shrinkage of nearby, but unstimulated spines. Interestingly, even when structural potentiation was switched off through inhibition of Ca^2+^/calmodulin-dependent protein kinase II (CaMKII), the heterosynaptic effect persists, suggesting a model in which spines send and receive shrinkage signals instead of competing for limited resources. Moreover, the observed dendritic locality is consistent with work on local or branch specific conservation of total synaptic conductance [106, 111].

Finally, in some cases heterosynaptic plasticity might not actually be heterosynaptic, as it may still depend on low levels of spontaneous synaptic activity in unstimulated synapses [112]. A recent biophysical model derived from synaptic plasticity data [113] actually requires such low-levels of activity to match the data. Interestingly, this model suggests such “heterosynaptic” plasticity could arise as a consequence of a rapid (timescale ∼ 12s) homeostatic sliding threshold possibly related to autophosphorylation of CaMKII.

While heterosynaptic plasticity, like synaptic scaling, can have a stabilizing effect, it is distinct from synaptic scaling in two ways. First, heterosynaptic plasticity need not multiplicatively scale all weights in the same manner. Second, it is unclear whether heterosynaptic plasticity in general drives neuronal activity variables to a specific set point, like synaptic scaling does [24]. Thus, while heterosynaptic plasticity has the rapidity to act as a putative RCP, its functional utility in storing memories requires further empirical and theoretical study, as it may lack the ability to precisely preserve ratios of synaptic strengths. However, its clear functional utility in preventing instability (Fig. 3d), along with even an approximate preservation in ratios of strengths, could potentially endow the interaction of heterosynaptic plasticity and Hebbian learning with the ability to stably learn and remember memories. Further network modeling, building on promising heterosynaptic plasticity models [30, 82, 113], could be highly instructive in elucidating the precise properties, beyond rapidity, a putative RCP must obey in order to provide appropriate competition and stability to Hebbian plasticity.

## Conclusion

The trinity of Hebbian plasticity, competition and stability are presumed to be crucial for effective learning and memory. However, a detailed theoretical and empirical understanding of how these diverse elements conspire to functionally shape neurobiological circuits is still missing. Here we have focused on one striking difference between existing models and neurobiology: the paradoxical separation of timescales between Hebbian and homeostatic plasticity. In models, such a separation of timescales typically leads to instability, unless plasticity is constrained by RCPs that act much faster than observed forms of homeostatic plasticity. In principle, RCPs could be implemented at various spatial scales. Here we have primarily discussed different forms of heterosynaptic plasticity and processes involving synaptic inhibition as possible candidates. However, rapid processes involving neuromodulation, glial interactions, or intrinsic plasticity [114–117] could also constitute RCPs. Thus, identifying the key neurobiological processes that provide stability and competition to Hebbian learning rules remains an important direction for future research.

## Acknowledgements

FZ was supported by the SNSF (Swiss National Science Foundation). SG was supported by the Burroughs Wellcome, Sloan, McKnight, Simons and James S. McDonnell foundations and the Office of Naval Research. WG was supported for this work by the European Research Council under grant agreement number 268689 (MultiRules) and by the European Community’s Seventh Framework Program under grant no. 604102 (Human Brain Project).

